# Regulation of ammonium acquisition and use in *Oryza longistaminata* ramets under nitrogen source heterogeneity

**DOI:** 10.1101/2021.08.24.457502

**Authors:** Misato Kawai, Ryo Tabata, Miwa Ohashi, Haruno Honda, Takehiro Kamiya, Mikiko Kojima, Yumiko Takebayashi, Shunsuke Oishi, Satoru Okamoto, Takushi Hachiya, Hitoshi Sakakibara

## Abstract

*Oryza longistaminata*, a wild rice, vegetatively reproduces and forms a networked clonal colony consisting of ramets connected by rhizomes. Although water, nutrients, and other molecules can be transferred between ramets via the rhizomes, inter-ramet communication in response to spatially heterogeneous nitrogen availability is not well understood. We studied the response of ramet pairs to heterogeneous nitrogen availability by using a split hydroponic system that allowed each ramet root to be exposed to different conditions. Ammonium uptake was compensatively enhanced in the sufficient-side root when roots of the ramet pairs were exposed to ammonium-sufficient and deficient conditions. Comparative transcriptome analysis revealed that a gene regulatory network for effective ammonium assimilation and amino acid biosynthesis was activated in the sufficient-side roots. Allocation of absorbed nitrogen from the nitrogen-sufficient to the deficient ramets was rather limited. Nitrogen was preferentially used for newly growing axillary buds on the sufficient-side ramets. Biosynthesis of *trans*-zeatin, a cytokinin, was up-regulated in response to the nitrogen supply, but *trans*-zeatin appears not to target the compensatory regulation. Our results also implied that the *O. longistaminata* ortholog of *OsCEP1* plays a role as a nitrogen-deficient signal in inter-ramet communication, providing compensatory up-regulation of nitrogen assimilatory genes. These results provide insights into the molecular basis for efficient growth strategies of asexually proliferating plants growing in areas where the distribution of ammonium ions is spatially heterogeneous.

**One sentence summary:** *Oryza longistaminata*, a rhizomatous wild rice, systemically regulates ammonium acquisition and use in response to spatially heterogeneous nitrogen availability via inter-ramet communication.

## Introduction

*Oryza longistaminata* is a wild rice species that preferentially grows in wetlands and vegetatively reproduces through rhizome growth that can be so vigorous as to occupy an area completely (Vaughan, 1994). Rhizome growth is characterized by the underground horizontal outgrowth of axillary buds to form new rhizomes that expand into new territory (Yoshida et al., 2016; Fan et al., 2017; Kyozuka, 2017; Bessho-Uehara et al., 2018; Toriba et al., 2020; Shibasaki et al., 2021). Rhizome tips developmentally transform into photosynthetic above-ground organs by growing into a new plantlet called a ramet. Continuous rhizome growth and subsequent transformation form a networked clonal colony. In addition to *O. longistaminata*, many other plant species vegetatively reproduce through rhizome growth, such as *Phyllostachys edulis* (Moso bamboo) and *Zoysia matrella* (Manila grass)(Guo et al., 2021), and show vigorous fertility. Since ramets are connected via rhizomes, water, nutrients, and other molecules can be transferred between ramets (De Kroon et al., 1998). Therefore, the growth and metabolism of a ramet are not totally independent of the neighboring ramets but have some influence on each other via the rhizomes.

Nitrogen is one of the most limiting macronutrients for plants, so nitrogen acquisition and use efficiency significantly affect plant growth and development. In submerged and reductive soil conditions where wetland rice grows, plants mainly use ammonium ions as the inorganic nitrogen source (Yoshida, 1981). In *Oryza sativa*, ammonium ions in the soil are taken up by ammonium transporters (AMTs)(Sonoda et al., 2003; Suenaga et al., 2003; Yuan et al., 2007; Li et al., 2016; Jia and von Wirén, 2020; Lee et al., 2020a; Konishi and Ma, 2021), and then are initially assimilated into glutamine and glutamate by the glutamine synthetase (GS)/ glutamate synthetase (GOGAT) cycle (Lea and Miflin, 1974). The cycle is mainly composed of cytosolic GS and plastidic NADH-GOGAT isoforms in the root (Ishiyama et al., 2004; Tabuchi et al., 2005; Tabuchi et al., 2007; Funayama et al., 2013; Yamaya and Kusano, 2014; Ji et al., 2019; Lee et al., 2020b). Several amino acids are synthesized by amino-transfer reactions from glutamine and glutamate to organic acids derived from glycolysis, the tricarboxylic acid cycle, and the pentose phosphate pathway. Previous studies using *O. sativa*, *Arabidopsis thaliana*, and *Nicotiana tabacum* revealed a wide and versatile regulatory network of gene expression in response to nitrogen (nitrate or ammonium) nutrition, including genes for the processes of uptake, assimilation, amino acid synthesis, carbon skeleton supply, and hormone signaling (Scheible et al., 1997; Scheible et al., 2004; Wang et al., 2004; Sakakibara et al., 2006; Chandran et al., 2016; Yang et al., 2017; Sun et al., 2020).

Plants optimize their growth and development according to the amount of available nitrogen, and signaling molecules, including phytohormones and peptide hormones, play a key role in this regulation (Nishida and Suzaki, 2018; Ruffel, 2018; Sakakibara, 2021; Wheeldon and Bennett, 2021). Cytokinin, especially *trans*-zeatin (tZ), promotes nitrogen-responsive shoot growth via modulating shoot meristem activity (Kiba et al., 2013; Davière and Achard, 2017; Kang et al., 2017; Osugi et al., 2017; Landrein et al., 2018). An abundant nitrate supply promotes *de novo* tZ biosynthesis via the up-regulation of *ADENOSINE PHOSPHATE-ISOPENTENYLTRANSFERASE3* (*IPT3*) and *CYP735A2* in Arabidopsis (Takei et al., 2004; Maeda et al., 2018; Naulin et al., 2019; Sakakibara, 2021). In *O. sativa*, a glutamine-related signal stimulates *OsIPT4* and *OsIPT5* expression that are involved in axillary bud outgrowth from shoots (Kamada-Nobusada et al., 2013; Ohashi et al., 2017). In *O. longistaminata*, the outgrowth of rhizome axillary buds in response to nitrogen nutrition is regulated by a similar process to that of the axillary shoot bud in *O. sativa*, despite differences in the physiological roles of these organs (Shibasaki et al., 2021). In contrast, strigolactone is involved in the nutrient-responsive suppression of shoot axillary bud outgrowth (Gomez-Roldan et al., 2008; Umehara et al., 2008; Minakuchi et al., 2010; Umehara et al., 2010). Strigolactone biosynthetic genes are known to be up-regulated in the roots of *O. sativa* and *Zea mays* under nitrogen-deficient (Sun et al., 2014; Xu et al., 2015; Ravazzolo et al., 2019; Bellegarde and Sakakibara, 2021; Ravazzolo et al., 2021) and phosphorus-deficient (Umehara et al., 2010) conditions.

It is essential for plants to efficiently acquire nutrients for optimal growth and development even though mineral nutrients are heterogeneously distributed in the soil. Recent studies using a split-root system in Arabidopsis showed that when part of the root system is subjected to nitrogen deficiency, C-TERMINALLY ENCODED PEPTIDE1 (CEP1), synthesized in the deficient root, is translocated to shoots via the xylem (Ohyama et al., 2008; Tabata et al., 2014). The perception of CEP1 by the CEP1 RECEPTOR (CEPR) in shoots triggers the expression of CEP DOWNSTREAM (CEPD) proteins that are translocated to the root system via the phloem, thereby promoting compensatory nitrogen uptake by inducing the expression of high-affinity nitrate transporters, including *NITRATE TRANSPORTER2.1* (*NRT2.1*) in the nitrate-ample side (Tabata et al., 2014; Okamoto et al., 2016; Ohkubo et al., 2017; Ruffel and Gojon, 2017; Ota et al., 2020; Ohkubo et al., 2021). In rhizomatous plants, ramets connected by rhizomes are regarded as one individual; however, it remains unknown whether each ramet responds to nitrogen availability independently or compensatively when growing in areas where nitrogen distribution is spatially heterogeneous.

In this study, we characterized the response of *O. longistaminata* ramet pairs, connected by a rhizome, to spatially heterogeneous nitrogen conditions and found a compensatory gene regulatory network for effective acquisition and assimilation of ammonium ions in the ample-side ramet root. Cytokinin biosynthesis was up-regulated in response to ammonium supply, but this increase appears to be independent of the compensatory regulation. Our results also imply that an *O. longistaminata* ortholog of the *OsCEP1* gene plays a role in inter-ramet communication.

This study provides valuable hints for understanding the molecular basis of efficient growth strategies of vegetatively proliferating plants.

## Results

### Compensatory promotion of ammonium uptake in response to a spatially heterogeneous ammonium supply in *O. longistaminata* ramet pairs

To examine nitrogen-related inter-ramet communication via the rhizome, we established a split-hydroponic experimental system for *O. longistaminata* ramet pairs (Supplemental Fig. S1). After preparing young ramet pairs of comparable growth stages that developed at adjacent rhizome nodes, we treated the roots of each ramet to different levels of nitrogen nutrition to mimic a condition of spatially heterogeneous nitrogen availability. We set up three conditions: (i) both ramet roots were exposed to 2.5 mM NH_4_Cl (+N), (ii) one ramet root was exposed to 2.5 mM NH_4_Cl (+N split), and the other was exposed to 0 mM NH_4_Cl (-N split), (iii) both ramet roots were exposed to 0 mM NH_4_Cl (-N) (Fig. 1A and Supplemental Fig. S1F). We compared the absorption activity of ammonium between the +N and +N split roots using ^15^N-labeled NH_4_Cl as the tracer. The results showed that the ammonium absorption activity was significantly increased by about 1.6-fold in the +N split ramets compared to the +N ramets (Fig. 1B). Expression of the *OlAMT1;2* and *OlAMT1;3* genes, orthologs of *O. sativa AMT1;2* and *AMT1;3*, that encode ammonium-inducible ammonium transporters, was approximately 1.8- and 3.6-fold higher in the roots of +N split than those of +N, respectively (Fig. 1C). These results suggest that when *O. longistaminata* ramet pairs are exposed to ammonium ions that are spatially heterogeneous, uptake is compensatively enhanced in the ramet exposed to the ammonium-rich condition and is accompanied by an up-regulation in *OlAMT1;2* and *OlAMT1;3* expression.

**Figure 1.**
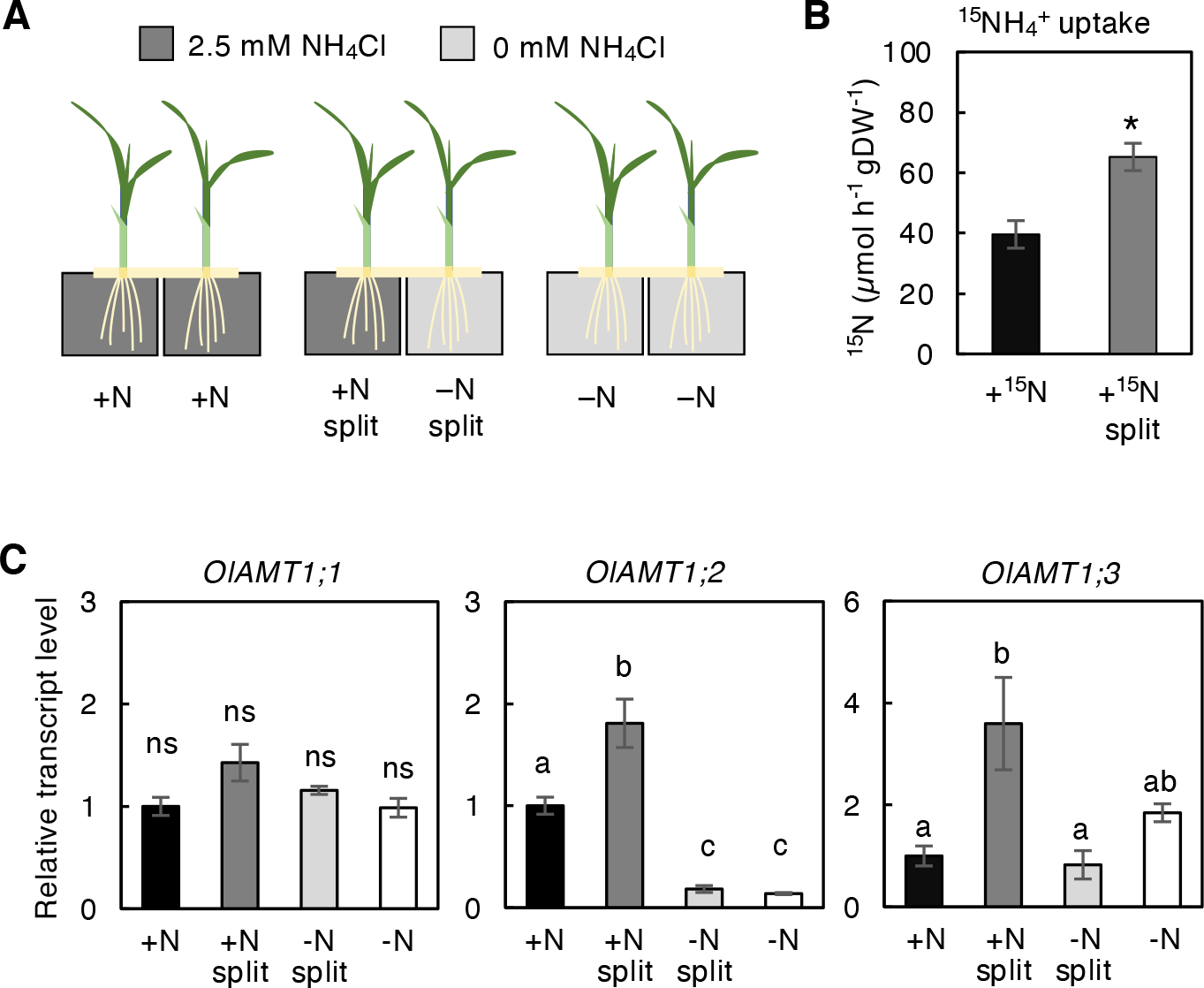
Compensatory up-regulation of ammonium ion uptake in response to spatially heterogeneous nitrogen availability. A, A schematic representation of the split hydroponic experiment system in which the roots of a ramet pair were separately exposed to independent nitrogen conditions. Details of the experimental design are shown in Fig. S1. B, Compensatory uptake of ammonium ion in +N split roots compared to +N. Ammonium ion uptake activity was measured using ^15^NH_4_^+^ as the tracer. Error bars represent SE of values for biological replicates (n = 3). \**p* < 0.05 (Student’s *t-*test). DW, dry weight. C, Accumulation pattern of *OlAMT1;1*, *OlAMT1;2*, and *OlAMT1;3* transcripts in the roots of *O. longistaminata* ramet pairs after a 24-h split treatment. Transcript abundance, normalized to *OlTBC*, is expressed relative to that of +N, defined as 1. Error bars represent SE of values for biological replicates (n = 3 or 4). Different lowercase letters at the top of each column denote statistically significant differences by Tukey’s honestly significant difference test (HSD) (*p* < 0.05). ns, not significant.

### Compensatory regulation of amino acid biosynthesis and related genes in response to a spatially heterogeneous ammonium supply

To explore the molecular basis underlying the compensatory response to a heterogeneous ammonium supply, we analyzed the transcriptome using RNA sequencing (RNAseq) of the roots at 24 h after the start of the +N, +N split, -N split, and -N treatments. Up- and down-regulated genes were identified from the expression data of the +N split, -N split, and -N treatments by comparison with that of the +N condition (false discovery rate [FDR] < 0.05). In comparison to the +N condition, 416 genes were significantly up-regulated in +N split root (Fig. 2A and Fig. 2B, Supplemental Table S1). Also, 2361 and 2776 genes were up-regulated in +N roots compared to the -N and -N split treatments, respectively (Fig. 2B, Supplemental Tables S2 and S3). Among the 416 genes, the identity of 295 genes overlapped with both the up-regulated genes in the +N treatment compared to the -N treatment and those in the +N treatment compared to the -N split treatment (110 genes), or only with those in the +N treatment compared to the -N split treatment (179 genes) and to the -N treatment (6 genes) (Fig. 2B). Gene Ontology (GO) analysis showed that genes associated with metabolic processes related to nitrogen and carbon, such as glycolysis (*p* = 2.7e-8), monosaccharide metabolic processes (*p* = 1.8e-7), oxoacid metabolic processes (*p* = 3.5e-7), and cellular amino acid metabolic processes (4.6e-6), were enriched among the 416 genes (Fig. 2C). Specifically, the up-regulated genes included those involved in nitrogen assimilation and amino acid synthesis, such as genes for a cytosolic glutamine synthetase *OlGS1;2*, an NADH-dependent glutamate synthase *OlNADH-GOGAT2*, an aspartate aminotransferase *OlAspAT*, an aspartate semialdehyde dehydrogenase *OlASADH*, an aspartate kinase-homoserine dehydrogenase *OlAK-HSDH1*, an alanine aminotransferase *OlAlaAT1*, and an aspartate kinase *OlAK* (Supplemental Table S1). Using reverse transcription-quantitative PCR (RT-qPCR), we confirmed that these genes were significantly up-regulated in +N split roots compared to +N (Fig. 3B). In addition, the expression of glycolytic enzyme genes involved in supplying carbon skeletons for amino acid biosynthesis, such as glucose-6-phosphate isomerase (*OlGPI*), triosephosphate isomerase (*OlTPI1*), fructose 1,6-bisphosphate aldolase (*OlALDC1*), glyceraldehyde 3-phosphate dehydrogenase (*OlGAPC1*), phosphofructokinase (*OlPFK5*), enolase (*OlENO1*), a glycolysis-related pyruvate orthophosphate dikinase (*OlPPDKA*), and phosphoenolpyruvate carboxylase (*OlPPC4*) were also significantly up-regulated in the +N split root condition compared to the +N treatment (Fig. 3A, Supplemental Table S1). These results suggest that a compensatory gene regulatory network to acquire and assimilate ammonium effectively is activated in the ramet roots on the nitrogen-sufficient side of the ramet pairs.

**Figure 2.**
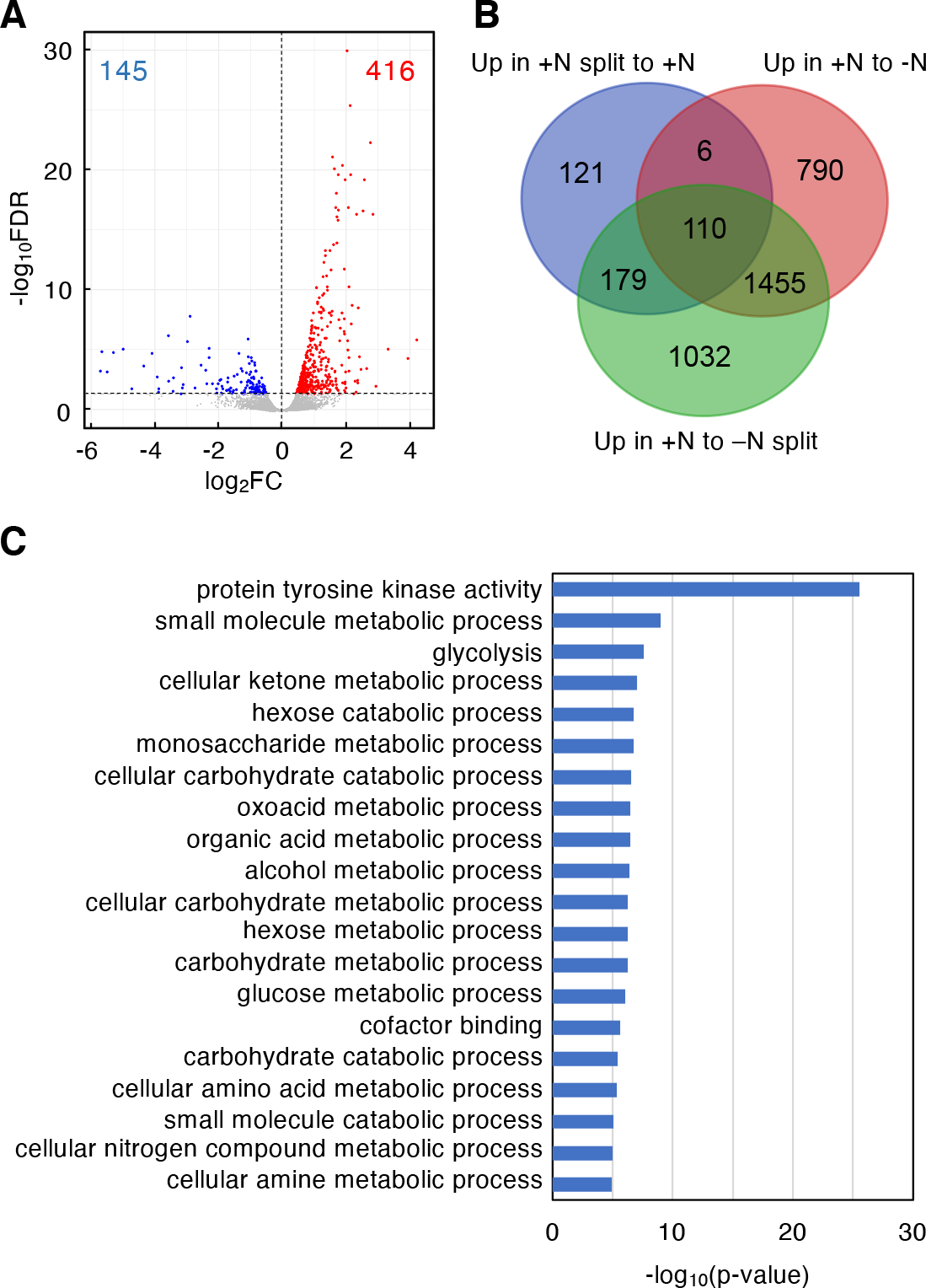
Transcriptome changes in the roots of ramet pairs in response to spatially heterogeneous nitrogen availability. A, A volcano plot showing differentially expressed genes (FDR < 0.05) between +N and +N split roots. Significantly up- regulated or down-regulated genes in roots from the +N split treatment are shown as red or blue dots, respectively. Numbers in red or blue indicate the total number of up-regulated or down-regulated genes. B, A Venn diagram showing the overlap among the genes up-regulated in the +N split treatment compared to the +N treatment (Up in +N split to +N), those up-regulated in the +N treatment compared to the -N treatment (Up in +N to -N), and those up-regulated in the +N treatment compared to the -N split treatment (Up in +N to -N split). C, The top 20 enriched GO categories among genes up-regulated in the +N split treatment compared to the +N treatment. Highly redundant GO categories were manually removed.

**Figure 3.**
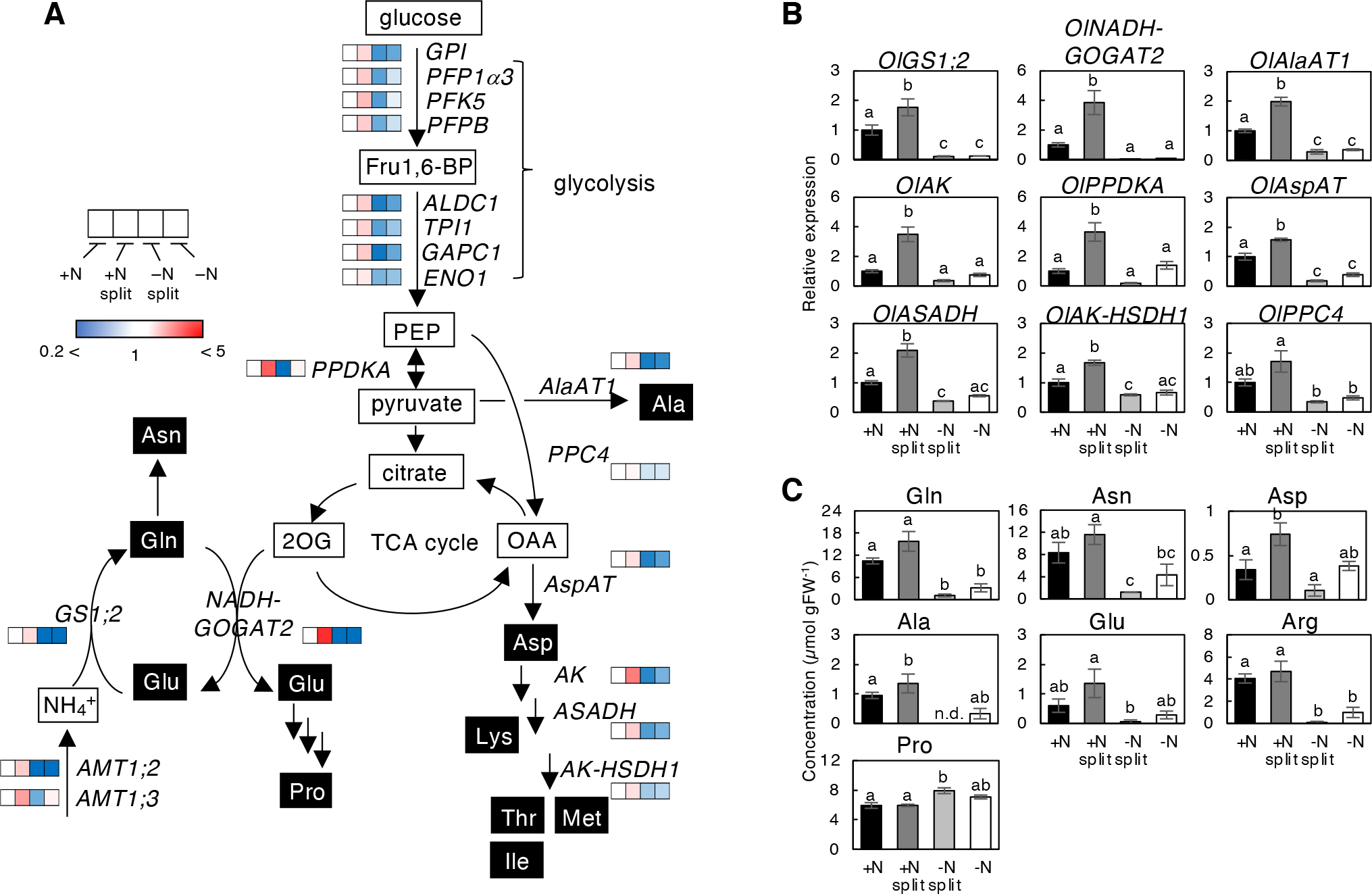
Metabolic responses to spatially heterogeneous nitrogen availability in the roots of *O. longistaminata* ramet pairs. A, The expression profile of genes involved in nitrogen assimilation, glycolysis and amino acid biosynthesis. The heatmaps in the pathways were deduced from RNAseq data. Fru1,6-BP, fructose 1,6-bisphosphate; PEP, phospho*enol-* pyruvate; OAA, oxaloacetate; 2OG. 2-oxoglutarate. The details of gene names are provided in Supplemental Table S1. B, Expression patterns of genes analyzed by RT-qPCR. The expression level of each gene, normalized to *OlTBC*, is expressed relative to that of the +N treatment defined as 1. C, Amino acid quantification. The concentrations of amino acids in the roots of each ramet pair after 48 h of split treatment are shown in histograms. Error bars represent the SE of values for biological replicates (n = 3 or 4 for RT-qPCR, and n = 4 for amino acid analysis). Different lowercase letters at the top of each column denote statistically significant differences by Tukey’s honestly significant difference test (HSD) (*p* < 0.05).

To know the impact of transcriptome changes on the accumulation level of amino acids, we analyzed the amino acid concentrations in roots from the +N, +N split, -N split, and -N treatments at a longer treatment period (48 h). As a result, the accumulation level of glutamine, asparagine, aspartate, alanine, glutamate, and arginine were significantly higher in the +N split roots than in the -N split roots (Fig. 3C). Among these amino acids, aspartate and alanine levels in roots from the +N split treatment were significantly higher than those in the +N treatment. Other amino acids except for arginine showed a similar tendency. These results are in line with the transcriptome changes in response to heterogeneous nitrogen availability.

On the other hand, 1848 and 2087 genes were up-regulated in the -N and -N split treatments compared to the +N condition, respectively (Supplemental Table S4 and S5). Overall, the expression of 1176 genes overlapped between the two conditions (Supplemental Fig. S2). One hundred and forty-five genes were up-regulated in the +N treatment compared to the +N split treatment (Supplemental Table S6), and 15 genes overlapped with the -N-up-regulated genes (Supplemental Fig. S2).

### Allocation of absorbed nitrogen between ramets via the rhizome when nitrogen availability is spatially heterogeneous

To examine the allocation of absorbed nitrogen in ramet pairs when nitrogen availability is spatially different, ^15^NH_4_Cl was fed to +N and +N split roots with the same conditions as shown in Fig. 1B, and the roots were grown for another 7 days with a non-labeled nitrogen source. In this experimental condition, ^15^N was detected in both the shoots and roots of the systemic +N and -N split ramets that had not been fed with ^15^NH_4_Cl (Fig. 4A), indicating that absorbed nitrogen is allocated to neighboring ramets via the rhizome. The percentage distribution of ^15^N was higher in the shoots than in the roots. Interestingly, the allocation of absorbed nitrogen from the +N split to the -N split ramets was less than that from the +N to the +N ramets.

**Figure 4.**
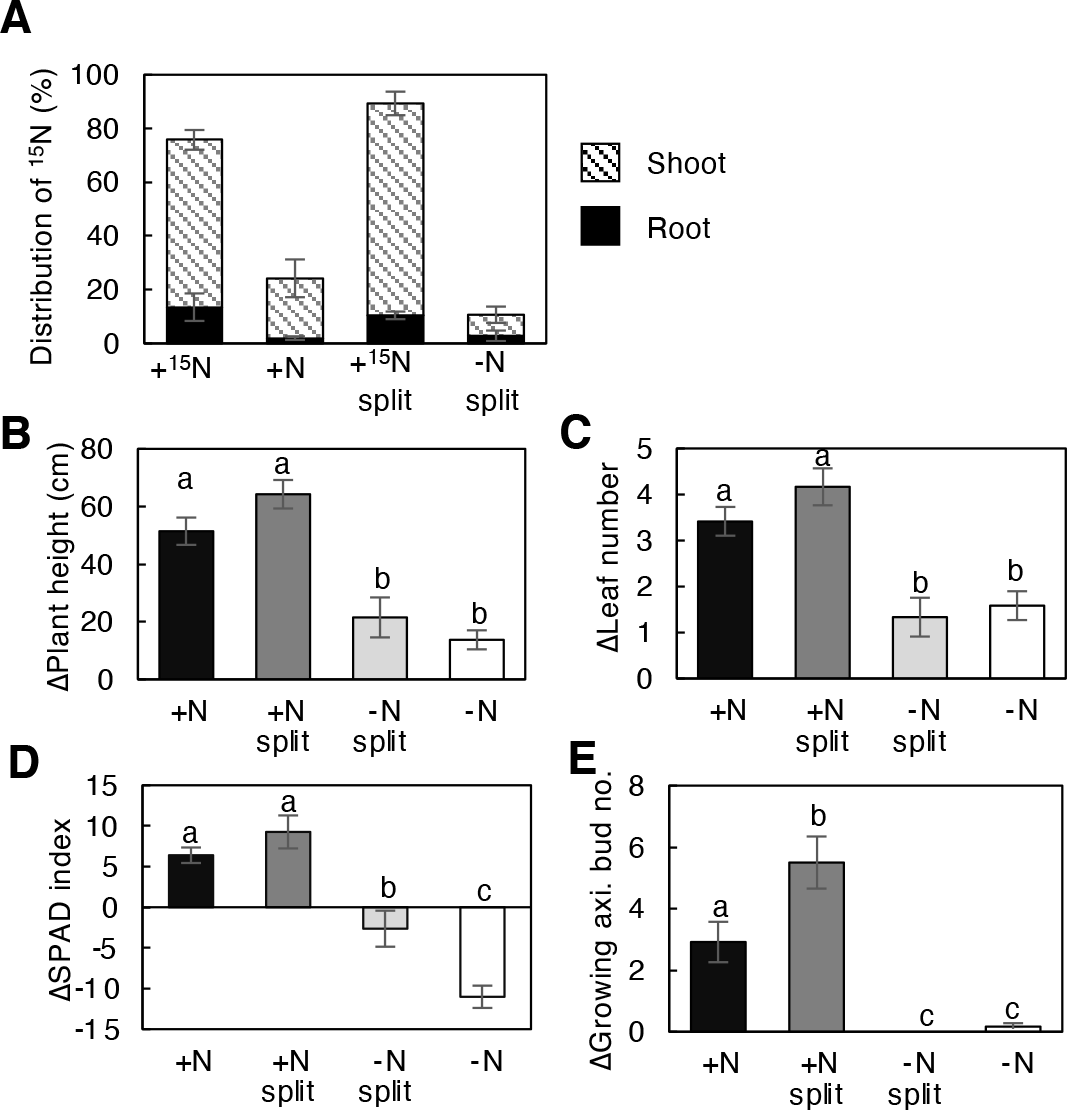
Allocation of absorbed nitrogen and growth profiles of ramet pairs. A, Distribution of ^15^N in roots and shoots 7 d after ^15^NH_4_Cl feeding. One of the ramets in each split-treated ramet pair was treated with 2.5 mM ^15^NH_4_Cl for 5 min and then returned to the normal split treatment. Error bars represent SE of values for biological replicates (n = 3). B to E, Changes in plant height (B, ΔPlant height), number of leaves (C, ΔLeaf number), chlorophyll content (D, ΔSPAD index), and number of growing axillary buds (E, ΔGrowing axl. bud no.) in each ramet after 5 weeks of split treatment. Error bars indicate the SE. Different lowercase letters at the top of each column denote statistically significant differences by Tukey’s honestly significant difference test (HSD) (*p* < 0.05). n = 6 to 12.

### Growth response of ramets in spatially heterogeneous nitrogen conditions

To know the long-term effects of nitrogen split treatment on the growth of ramets, we exposed ramet pairs to different nitrogen conditions for 5 weeks, and growth parameters for each ramet were monitored (Figs. 4B to E). Plant height, the number of fully developed leaves, chlorophyll content, and the number of growing axillary buds were significantly higher in the +N and +N split ramets than in the -N split and -N ramets. The chlorophyll content was lower in the -N split and -N ramet shoots, but the decrease was significantly alleviated in the -N split ramets compared to the -N ramets (Fig. 4D), suggesting that the allocated nitrogen was used to retain photosynthetic function. In contrast, the change in the number of growing axillary buds was almost zero for the -N and -N split ramets, whereas the axillary bud number increased significantly in the +N split and +N ramets (Fig. 4E). In particular, the axillary bud number was significantly higher in the +N split ramets than in the +N ramets. These results suggest that when neighboring ramets are exposed to different levels of nitrogen availability, the allocation of absorbed nitrogen from the N-sufficient ramet to the deficient ramet is rather limited and, nitrogen is preferentially allocated for newly growing axillary buds.

### Response of cytokinin and strigolactone biosynthetic genes in response to a heterogeneous nitrogen supply

To gain insight into the growth promotion of axillary buds in the +N split ramets, we focused on cytokinin and strigolactone, two phytohormone families that promote and inhibit axillary bud outgrowth, respectively. We analyzed the expression levels of cytokinin and strigolactone biosynthetic genes by RT-qPCR using the roots of ramet pairs that had been split-treated for 7 days. The expression levels of cytokinin biosynthetic genes, *OlIPT4*, *OlIPT5*, *OlCYP735A3*, and *OlCYP735A4* were higher in the roots of the +N and +N split ramets than in those of the -N and -N split ramets (Fig. 5A), and there was no difference between the +N and +N split conditions.

**Figure 5.**
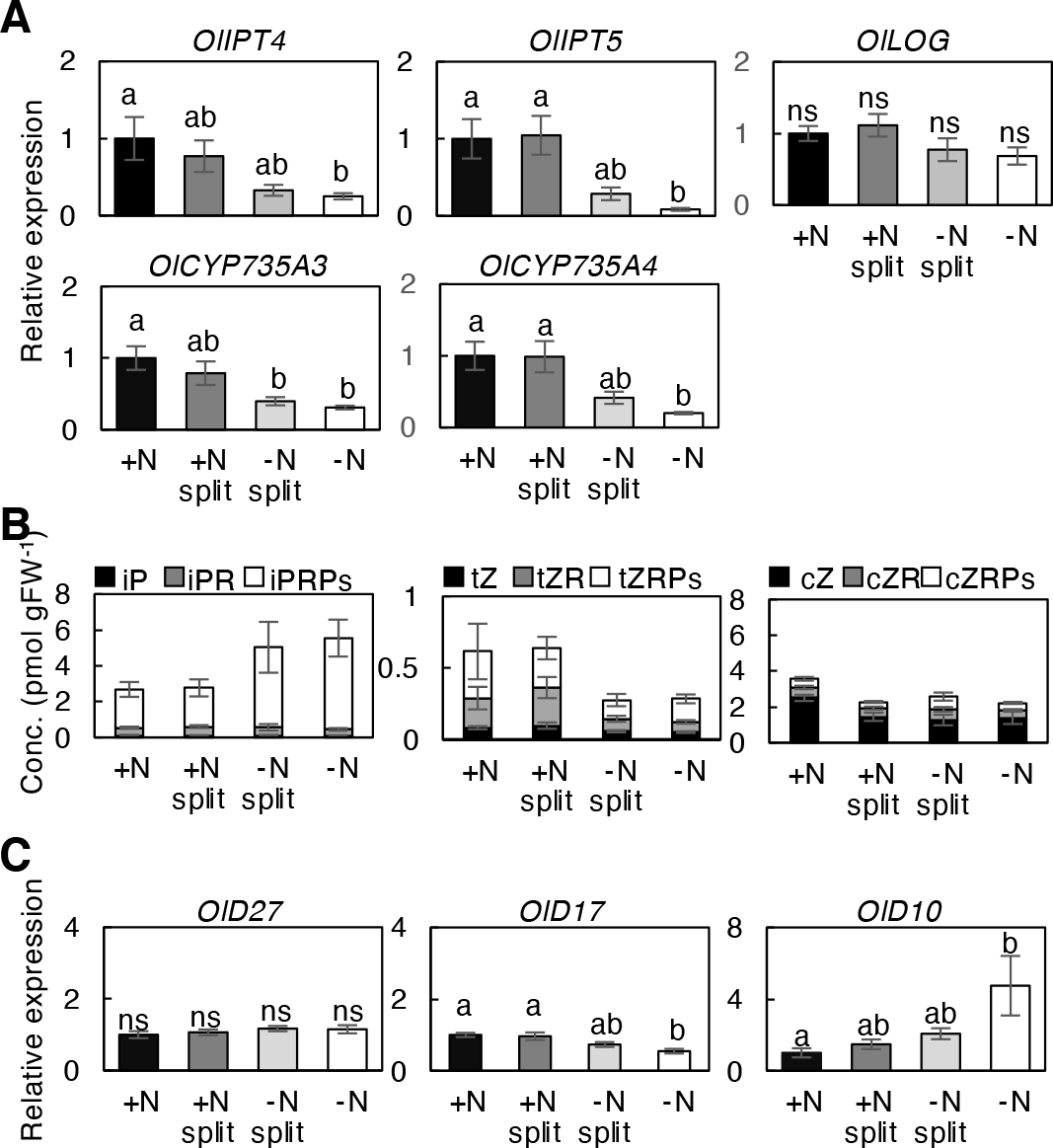
Response of cytokinin and strigolactone biosynthesis to heterogeneous nitrogen conditions. A, Relative expression level of cytokinin biosynthetic genes in ramet roots in response to different nitrogen treatments. Transcript abundance in roots of each ramet after 1 week of the split treatment was analyzed by RT-qPCR. B, Cytokinin concentration in the roots of each ramet after 1 week of the split treatment. The complete data set is presented in Supplemental Table S7. tZ, *trans*-zeatin; tZR, tZ riboside; tZRPs, tZR 5’-phosphates; cZ, *cis*-zeatin; cZR, cZ riboside; cZRPs, cZR 5’-phosphates; iP, *N6*-(Δ2-isopentenyl)adenine; iPR, iP riboside; iPRPs, iPR 5’-phosphates; and FW, fresh weight. C, Transcript abundance of strigolactone biosynthetic genes in ramet roots in response to different nitrogen conditions. In (A) and (C), the expression level of each gene normalized by *TBC* is expressed relative to that of the +N treatment defined as 1. Error bars represent the SE of values for biological replicates (n = 4). Different lowercase letters at the top of each column denote statistically significant differences by Tukey’s honestly significant difference test (HSD) (*p* < 0.05).

We also analyzed the concentration of cytokinins in the roots. The level of *trans*-zeatin (tZ)-type cytokinins, including the riboside and ribotide precursors, was higher in the +N and +N split roots compared to the -N split and -N roots, whereas the levels of *N^6^*-(Δ^2^-isopentenyl)adenine (iP)-type cytokinins were somewhat lower (Fig. 5B, Supplemental Table S7). The tZ-type cytokinin content in the +N and +N split roots was comparable, suggesting that de novo tZ-type cytokinin biosynthesis is enhanced in response to the ammonium supply but is not under compensatory regulation.

Although strigolactone species in *O. longistaminata* have not been well characterized yet, we analyzed orthologs of *O. sativa D27*, *D17*, and *D10* (*OlD27*, *OlD17*, and *OlD10*, respectively) encoding enzymes involved in the production of carlactone, an intermediate of strigolactone biosynthesis (Alder et al., 2012). The expression level of *OlD10* was significantly higher in -N ramet roots than in other conditions, although *OlD27* and *OlD17* expression levels were essentially similar and slightly lower, respectively (Fig. 5C). This result implies that de novo strigolactone biosynthesis might be up-regulated in nitrogen-deficient roots.

### Possible involvement of a CEP1-type peptide in inter-ramet nitrogen-deficiency signaling

In Arabidopsis, CEP1 plays a key role in the systemic regulation of nitrate acquisition in response to heterogeneous nitrogen conditions as a root-to-shoot signaling molecule (Tabata et al., 2014; Okamoto et al., 2016). To investigate whether a CEP1-type peptide is involved in the observed inter-ramet communication, we analyzed the response of *O. longistaminata* CEP gene orthologs to spatially heterogeneous nitrogen availability. In *O. sativa*, 15 genes encode CEPs (Sui et al., 2016). We searched for the orthologs in *O. longistaminata* and found 15 sequences corresponding to each of the *O. sativa* genes (Supplemental Table S8), and eight of the genes (*OlCEP5*, *OlCEP6.1*, *OlCEP9*, *OlCEP10*, *OlCEP11*, *OlCEP12*, *OlCEP14*, and *OlCEP15*) were annotated in our RNAseq data. However, expression of these eight genes was not significantly different in -N roots compared to +N roots. Next, we focused on *OlCEP1* and analyzed its expression level by RT-qPCR because the peptide sequences encoded by *OsCEP1* (also *OlCEP1*) belong to the same group (group I) as Arabidopsis CEP1 based on the structural features (Delay et al., 2013; Sui et al., 2016). In the RT-qPCR analysis, the expression level of *OlCEP1* in -N roots was significantly higher than in the +N and +N split roots (Fig. 6A).

**Figure 6.**
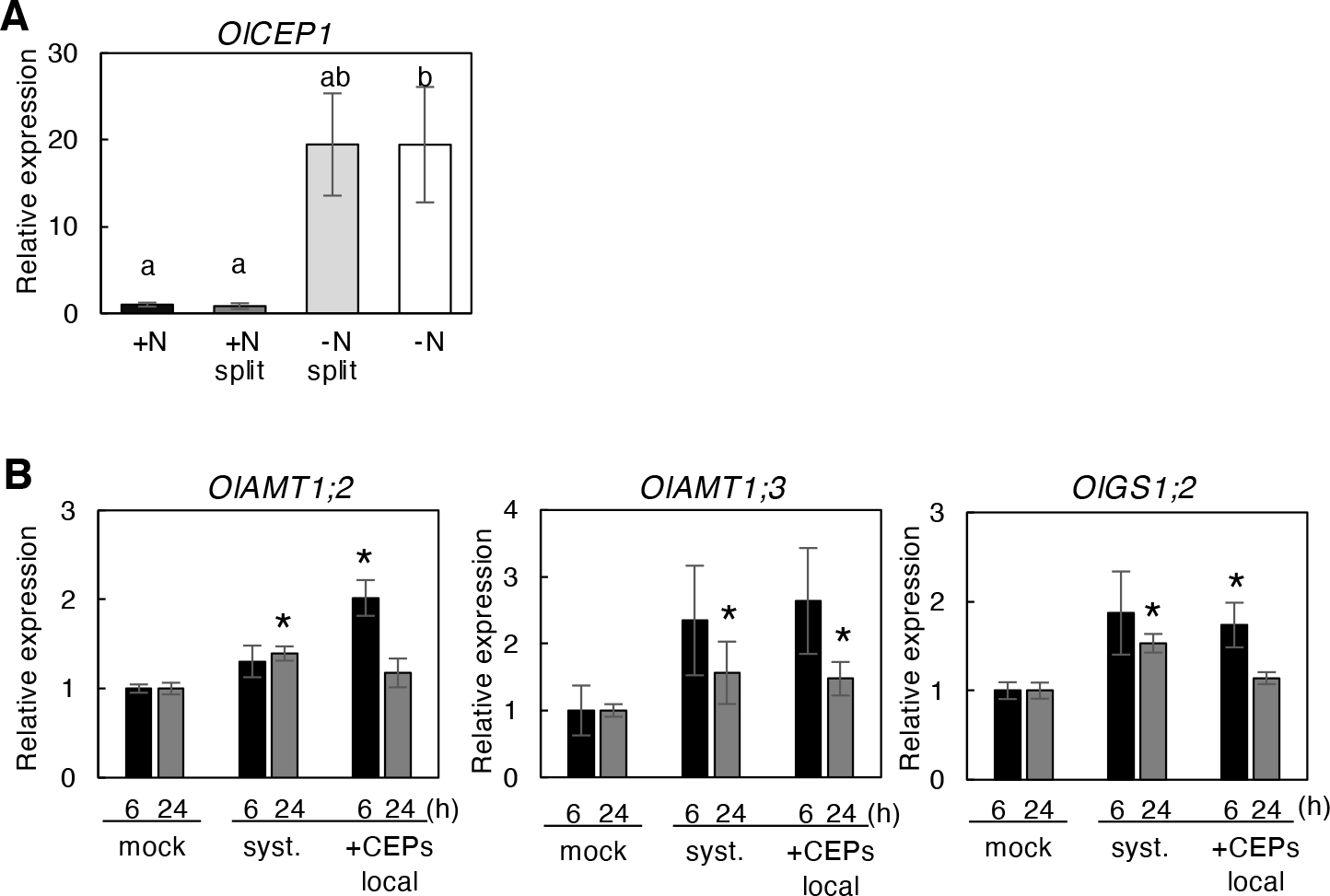
A, Expression pattern of the *O. longistaminata* ortholog of *OsCEP1* (*OlCEP1*) in the roots of ramet pairs after a 24-h split treatment. The expression level of each gene normalized by *TBC* is expressed relative to that of the +N treatment defined as 1. Error bars represent SE of values for biological replicates (n = 3 or 4). Different lowercase letters at the top of each column denote statistically significant differences by Tukey’s honestly significant difference test (HSD) (*p* < 0.05). B, Expression of *OlAMT1;2, OlAMT1;3,* and *OlGS1;2* genes in the ramet roots in response to exogenously supplied CEP1 peptides. The CEP1 peptide mix was applied to the roots of one of the ramet pairs, and gene expression in the local (+CEP local) and systemic side (systemic) roots after 6 and 24 h treatment was measured. The expression level of each gene, normalized by *TBC*, is expressed relative to that of the +N mock treatment defined as 1. Error bars represent SE of values for biological replicates (n = 3 or 4). \**p* < 0.05 (Student’s t-test) compared to the corresponding mock treatment.

To gain further insight into the involvement of *OlCEP1* gene products in nitrogen-related ramet-to-ramet signaling, we examined the effect of an exogenous application of OlCEP1 peptides on the expression of *OlAMT1;2, OlAMT1;3,* and *OlGS1;2* genes. In addition to *OsCEP1*, the coding sequence of the *OlCEP1* gene contains multiple CEP sequences containing proline residues. Thus, we synthesized peptides OlCEP1a to OlCEP1d (See Materials and Methods) and used a mixture. We treated ramet roots with the peptide mixture under nitrogen-sufficient conditions to repress the endogenous *OlCEP1* expression. When we applied OlCEP1 peptides to the roots of one side of the ramets, the accumulation level of transcripts for *OlAMT1;2, OlAMT1;3,* and *OlGS1;2* increased in both the local side and systemic side of the ramet roots in 6 h. The transcript level significance was maintained in the systemic side of the root even at 24 h after treatment (Fig. 6B), suggesting that CEP1-related signaling is possibly involved in the systemic regulation of the genes via the rhizomes.

We further explored homologs of the *CEPR* and *CEPD* genes in *O. longistaminata*. Since no previous studies of CEPR and CEPD have been conducted in *O. sativa* and *O. longistminata*, we identified *Ol12G001732* as the closest homolog of Arabidopsis CEPR1 in the PLAZA database (Van Bel et al., 2018) and ten CC-type glutaredoxin genes (GRX) (Garg et al., 2010) as the most closely related homologs of Arabidopsis CEPD (Supplemental Figs. S4 and S5; Supplemental Table S8). Our RT-qPCR analysis showed that the expression of *Ol12G001732* was highest in the +N split shoot (Supplemental Fig. S6A). Up-regulation of *Ol12G001732* was also observed in the CEP1 peptide treatment experiment (Supplemental Fig. S7), suggesting that the CEPR homolog might play a role in ramet-to-ramet nitrogen signaling in *O. longistaminata*. On the other hand, there was no significant difference in the expression of the *GRXs* in the spatially heterogeneous nitrogen condition except for *OlGRX15*, but the up-regulation of *OlGRX15* expression was not +N split specific (Supplemental Fig. S6B).

## Discussion

In this study, we demonstrated inter-ramet communication occurs in *O. longistaminata* via rhizomes for a systemic response to spatially heterogeneous nitrogen availability. When ramet pairs of *O. longistaminata* connected by a rhizome were exposed to different ammonium ion conditions, a series of gene networks that allow complementary absorption and assimilation of ammonium was activated in the root of the ramet in the nitrogen-sufficient condition. This network also included genes capable of supplying the carbon skeletons for amino acid synthesis, such as those involved in glycolysis. The expression level of these genes in the nitrogen-sufficient side of the heterogeneous condition was higher than in the nitrogen homogeneously sufficient condition, suggesting that the nitrogen-deficient side ramet conveyed some kind of nitrogen deficiency signal to the sufficient side ramet via the rhizome to trigger the systemic response.

In our transcriptome analysis, a large part of the compensatively up-regulated genes overlapped with ammonium-responsive genes (295 genes/416 genes, Fig. 2B), indicating that the expression of the compensatory genes is up-regulated by an ammonium-signal *per se* but further boosted by the input of a systemic nitrogen-deficient signal from the adjacent ramet. It is not clear at present whether a de-repression or a more facilitative regulatory event underlies the process.

A small part of the nitrogen absorbed in the ramet roots in the heterogeneous nitrogen-sufficient condition (+N split) was distributed to the adjacent nitrogen-deficient ramets (-N split), contributing to the maintenance of the chlorophyll content. However, no evidence was obtained to suggest that the distributed nitrogen was used for growth of the nitrogen-deficient side ramet. In a previous study in *Carex flacca* examining water and nitrogen transfer between ramets via the rhizome, the direction of nitrogen nutrient transport depended on the direction of water transport (De Kroon et al., 1998). In rice cultivars, the nitrogen-sufficient condition elevates the leaf transpiration rate compared to the limited condition (Xiong et al., 2015). Thus, it is likely that the translocation of nitrogen from the sufficient side to the deficient side against the water flow is limited between rhizome-connected ramets experiencing different nitrogen conditions.

Our results suggest that most compensatively acquired nitrogen is locally used for growth on the sufficient side. Given that an *O. longistaminata* cluster is a single clonal colony, this use of nitrogen might be a strategy to ensure the colony’s survival in limited and heterogeneous nitrogen conditions.

Notably, genes for the biosynthesis of tZ-type cytokinins, *IPTs* and *CYP735As*, were up-regulated by the ammonium supply. Still, the expression levels and accumulation of tZ-type cytokinins (tZ and its precursors) were comparable between the +N and +N split treatments in our experiments (Fig. 5B). Therefore, we hypothesize that cytokinin synthesis is similarly up-regulated in response to both homogeneous and heterogeneous nitrogen conditions.

Ammonium acquisition and amino acid synthesis, however, are compensatively enhanced in the nitrogen-rich ramet roots under heterogeneous conditions, resulting in a greater supply of nitrogen assimilates for axillary bud outgrowth. Both the cytokinin signal and the supply of building blocks could contribute to the preferential axillary shoot growth.

In our analysis, iP and its precursors were in low abundance in the +N and +N split roots, mainly due to the decreased levels of its ribotide precursor (iPRPs, Fig. 5B). At present, we do not have a clear explanation for the opposite trend despite the up-regulation of its biosynthetic genes, *OlIPTs*. The iP ribotide precursor might be over consumed by CYP735As to produce tZ-type species.

Previous studies in *O. sativa* and *O. longistaminata* indicated that *NADH-GOGAT1* is under the same control as *IPT4* in the local nitrogen response and that glutamine-related signaling is involved in the regulation (Kamada-Nobusada et al., 2013; Ohashi et al., 2017; Shibasaki et al., 2021). *OlNADH-GOGAT2*, an isogene, is compensatively up-regulated in our experimental conditions, whereas *OlNADH-GOGAT1* had a similar expression pattern to *OlIPT4* (Supplemental Fig. S3). In *O. sativa*, *NADH-GOGAT2* is mainly expressed in mature leaves and plays a role in providing glutamate for the GS1;1 reaction in vascular tissues for nitrogen remobilization and recycling (Tabuchi et al., 2005; Tabuchi et al., 2007; Yamaya and Kusano, 2014). Our RNAseq data show that *OlGS1;1* is up-regulated in nitrogen-deficient conditions (Supplemental Tables S4 and S5) in a manner distinct from *OlNADH-GOGAT2*. Thus, the physiological role of the cytosolic GS isoforms and NADH-GOGAT isoforms might be different between *O. sativa* and *O. longistaminata*.

Systemic regulation of nitrate acquisition in response to heterogeneous nitrogen nutrient conditions by CEP1-CEPR-CEPD has been identified in Arabidopsis (Tabata et al., 2014; Okamoto et al., 2016; Ohkubo et al., 2017; Ruffel and Gojon, 2017; Ota et al., 2020; Ohkubo et al., 2021). In our analysis, the *O. longistaminata* ortholog of *OsCEP1* was markedly induced by nitrogen deprivation, and exogenous application of the synthetic CEP1 peptides increased the expression level of *OlAMT1;2, OlAMT1;3,* and *OlGS1;2* on the systemic side as well as the local side root. In addition, *Ol12G001732*, the closest ortholog of Arabidopsis *CEPR1*, was up-regulated in systemic side shoots in response to a spatially heterogeneous ammonium supply (Supplemental Fig. S6A) and also in systemic and local side shoots treated with CEP1 peptides (Supplemental Fig. S7). This finding implies that CEP plays a role in inter-ramet communication as a nitrogen-deficient signaling molecule via the rhizome. Up-regulation of *Ol12G001732* expression might sensitize the root-derived CEP1-signal. In contrast, there were no *GRX* genes whose expression increased only in +N split shoots in our experimental conditions (Supplemental Fig. S6B). At present, it is unclear whether the whole set of CEP-CPER-CEPD modules function in Oryza species. The systemic response mechanism responding to heterogeneous nitrogen conditions in monocots, including rice, is still largely unexplored.

Characterizing the whole system at the molecular level will require identifying and characterizing the CEP receptor and associated downstream factors in Oryza species.

## Materials and methods

### Plant materials and growth conditions

The perennial wild rice species *O. longistaminata* (IRGC10404) was hydroponically grown under natural light conditions in a greenhouse with a nutrient solution described by Kamachi et al. (1991), except that the solution contained 1 mM NH_4_Cl as the sole nitrogen source and was renewed once every 3 or 4 days. Pairs of young ramets grown on the proximal nodes of rhizomes were excised and used in experiments. The growth stage of all ramet pairs was similar; each ramet had two to three fully developed leaves (Supplemental Fig. S1).

### Split treatment

The roots of each ramet pair, connected by rhizomes, were incubated in two separate pots (Supplemental Fig. S1). One L of nutrient solution was used for each ramet. For the ±N split treatment, one of the ramet pairs was treated with the nutrient solution containing 2.5 mM NH_4_Cl (+N split), and the other was treated with 0 mM NH_4_Cl (-N split). For comparison, both ramet pairs were treated with 2.5 mM NH_4_Cl (+N) or 0 mM NH_4_Cl (-N). Treatments for the growth analysis, and cytokinin and strigolactone response analysis were conducted in a greenhouse. All other treatments were conducted in a growth cabinet (LPH-411PFQDT-SPC, NK System, Osaka, Japan) with the following environmental conditions: 16 h (28°C) light/8 h (24°C) dark at 400 µmol photons m^-2^ s^-1^.

### 15NH4 tracer experiments

The ramet pairs were initially incubated with water for 3 days, followed by a 24-h split treatment with stable isotope-free nitrogen nutrient solutions (2.5 mM NH_4_Cl for the +N ramet and the +N split ramet, and 0 mM NH_4_Cl for the -N ramet). Roots of the ramets were soaked in 800 mL of 1 mM CaSO_4_ for 1 min to remove the nitrogen treatment solution. The solution for the +N-treated side was then replaced with a solution containing 2.5 mM ^15^NH_4_Cl (^15^N 99 atom%, Shoko Science Co., Ltd., Yokohama, Japan), and ^15^NH_4_^+^ was allowed to be absorbed for 5 min. The roots of ramet pairs were then soaked in 600 mL of 1 mM CaSO_4_ for 1 min to wash out ^15^N and then treated again with the stable isotope-free +N or -N hydroponic solutions for 22 h to measure the ammonium absorption activity and for 7 days to measure the absorbed nitrogen distribution with daily changing of the culture solution in a growth cabinet (LPH-411PFQDT-SPC, NK System) with 16 h (30°C) light/8 h (30°C) dark at 400 µmol photons m^-2^ s^-1^.

The above-ground tissues, rhizomes, and roots were separately harvested and dried in an oven at 70°C for at least 5 days. All dried tissues were weighed and ground into fine powders. The ^15^N and total nitrogen contents were analyzed by Shoko Science Co. with an elemental analyzer-isotope ratio mass spectrometer (Flash2000-DELTA plus Advantage ConFlo System,

Thermo Fisher Scientific, Waltham, MA, United States).

The absorption activity of ammonium was calculated as follows. The increase in ^15^N (µmol) in each sample (^15^N_s_) was calculated from equation (1). W_s_, dry weight of the sample (gDW); N_s_, the total nitrogen concentration of the sample (%); ^15^N_s_, the ^15^N concentration of the sample; and ^15^N_0_, the naturally occurring ^15^N concentration (%) of the sample.

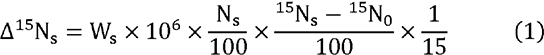

The Δ ^15^N of each sample (above-ground tissues and roots) were summed to obtain the increase of ^15^N in the whole ramet pair (Δ^15^N ). The absorption activity of ammonium (µmol gDW^-1^ h^-1^) in roots under +N or +N split conditions was calculated using equation (2). A, the absorption activity of ammonium; W_r_, the dry weight of the root that absorbed ^15^N.

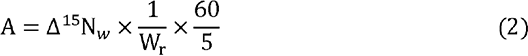

The increases in ^15^N_s_ in each part of the ramet pair (Δ^15^N_s_) were summed to obtain the increase of ^15^N in the whole ramet pair (Δ^15^N_w_), and the percentage (%) of ^15^N distributed in each part of the ramet pair was calculated using equation (3). P, the percentage of ^15^N distributed in each part.

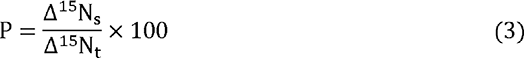

### Reverse transcription-quantitative PCR analysis

Total RNA was extracted from frozen and ground tissues using the RNeasy Plant Mini Kit (Qiagen, Hilden, Germany) with the RNase-Free-DNase Set (Qiagen) according to the supplier’s protocols. One µg of total RNA was used to synthesize cDNA using the ReverTra Ace qPCR Master Mix (TOYOBO) according to the supplier’s protocol. Twenty-ng cDNA was used for each qPCR reaction with the KAPA SYBR FAST qPCR Master Mix (2X) (KAPA Biosystems, Wilmington, MA, United States) and a real-time PCR system (Applied Biosystems QuantStudio 3). Expression levels were estimated using the relative quantification method (Livak and Schmittgen, 2001) with *OlTBC*, a homolog of TBC1 domain family member 22A (Maksup et al., 2013), the internal standard for normalization. Gene locus IDs and the specific primers used for amplification are listed in Supplemental Table S9.

### Determination of free amino acids

Free amino acids were extracted as described by Konishi et al. (2014). Derivatization of amino acids was carried out using the AccQ-Tag Ultra Derivatization Kit (Waters Corp., Milford, MA, United States). The resulting AccQ-Tag-labeled derivatives were separated and quantified using an HPLC System (Alliance 2695 HPLC system/2475, Waters Corp.) with the AccQTag Column (3.9×150 mm, Waters Corp.) as described in the instruction manual.

### RNA-Seq analysis

Libraries were prepared with the TruSeq Standard mRNA Library Prep Kit (Illumina, San Diego, CA, United States) using 1 µg of total RNA. Sequence analysis of 40 M reads was performed with a NovaSeq 6000 Sequencing System (Illumina). Library preparation and sequencing were conducted by Macrogen Japan, Inc. (Tokyo). The sequencing reads were mapped to the *O. longistaminata* genome obtained from the *Oryza longistaminata* Information Resource (http://olinfres.nig.ac.jp/)(Reuscher et al., 2018) using HISAT2 (Kim et al., 2019), followed by featureCounts (Liao et al., 2014) for counting reads and edgeR (Robinson et al., 2009) for differential expression analysis. Low expression genes were removed using filterByExpr function of edgeR with the default setting. We obtained the presumed function and MAPMAN BIN code for each gene and the presumed corresponding gene ID of *O. sativa* from Supplemental data 2 of Reuscher et al. (2018). Gene ontology (GO) analysis was performed using the corresponding *O. sativa* gene ID list with the analysis tool agriGO (http://systemsbiology.cau.edu.cn/agriGOv2/) (Du et al., 2010; Tian et al., 2017).

### Growth analysis

Ramet pairs at a similar growth stage were first grown hydroponically in water for 4 days and subsequently exposed to nitrogen split conditions for 5 weeks in a greenhouse using the hydroponic culture solution with (+N split, 2.5 mM NH_4_Cl) or without (-N split, 0 mM NH_4_Cl) nitrogen. The hydroponic solution was renewed every 3 to 4 days. Growth changes were analyzed at 7-day intervals by measuring plant height, the number of fully expanded leaves, the number of axillary buds growing more than 1 cm, and the chlorophyll content. The chlorophyll content (SPAD value) was measured with a SPAD-502 Plus Chlorophyll Meter (Konica Minolta, Tokyo, Japan) at the tip, middle, and basal parts of leaves, and the average was taken as the SPAD value. For the controls, both ramets of each pair were fed 2.5 mM NH_4_Cl (+N) or 0 mM NH_4_Cl (-N)-containing hydroponic solution for 5 weeks.

### Phytohormone quantification

Cytokinins were extracted and semi-purified from about 100 mg fresh weight of root tissues as described previously (Kojima et al., 2009). Cytokinins were quantified using an ultra-performance liquid chromatography (UPLC)-tandem quadrupole mass spectrometer (ACQUITY UPLC System/XEVO-TQXS; Waters Corp.) with an octadecylsilyl (ODS) column (ACQUITY UPLC HSS T3, 1.8 µm, 2.1 mm × 100 mm, Waters Corp.; Kojima et al., 2009).

### Peptide synthesis

OsCEP1a (DVRHypTNPGHSHypGIGH), OsCEP1b (DVRHypTNHypGHSHypGIGH), OsCEP1c (GVRHypTNPGHSHypGIGH), and OsCEP1d (GVRHypTNHypGHSHypGIGH) were synthesized on a CS 136X synthesizer (CS Bio, Menlo Park, CA, United States) using Fmoc solid phase peptide synthesis chemistry. Hydroxyproline (Hyp) was introduced with Fmoc-Hyp(Boc)-OH purchased from Watanabe Chemical Industries, Ltd. (Hiroshima, Japan). The obtained crude peptides were purified by reverse-phase HPLC on Jasco, Inc. (Tokyo, Japan) preparative instruments with a Jupiter C18 column (5 µm, 300 Å pore size, 21.2 mm internal diameter × 250 mm) (Phenomenex, Torrance, CA, United States), and lyophilized to yield pure peptides.

### Treatment with synthesized CEP1 peptides

Ramet pairs at a similar growth stage were first grown in hydroponic solution for 3 days in a growth chamber (LPH-411PFQDT-SPC, NK System). Some of the root tips (about 3 cm) from both ramets were removed with a razor blade to facilitate uptake of the exogenously supplied CEP1 peptides. Water on the root surface was wiped off with a Kim Towel (Nippon Paper Cresia, Tokyo, Japan). The root from one side of the ramet was immediately submerged into 100 mL of an ammonium solution containing CEP peptides (+CEPs local: 2.5 mM NH_4_Cl, 30 µM of each CEP1a-d). Simultaneously, the root on the other side was submerged in the ammonium solution (systemic: 2.5 mM NH_4_Cl) for the indicated period. For the mock control, the roots of each ramet pair were treated with 2.5 mM NH_4_Cl.

### Statistical analysis

Graphs represent the means ± SE of biological-independent experiments. The statistical significance was assayed using means a two-tailed Student’s *t-*test or Tukey’s honestly significant difference test (*p* < 0.05). The choice of test and the replicate numbers are provided in the corresponding figure legend.

### Accession numbers

RNA-seq data were deposited to the Gene Expression Omnibus (https://www.ncbi.nlm.nih.gov/geo/) under accession number GSE182486.

## Supplemental data

The following supplemental materials are available.

Supplemental Figure S1. The split hydroponic experimental system for *O. longistaminata* ramet pairs.

Supplemental Figure S2. A Venn diagram showing the overlap among genes up-regulated in the -N treatment compared to the +N treatment, those up-regulated in the -N split treatment compared to +N treatment, and those up-regulated in the +N treatment compared to +N split treatment.

Supplemental Figure S3. Transcript abundance of *OlNADH-GOGAT1* in the roots of ramet pairs after a 24-h split treatment as measured by RT-qPCR.

Supplemental Figure S4. Phylogenetic analysis of Arabidopsis CEPDs (AtCEPD1, AtCEPD2, AtCEDL1, AtCEPDL2) and *O. sativa* glutaredoxin family proteins, GRX and GRL. **Supplemental Figure S5**. An alignment of the amino acid sequences used for the phylogenetic analysis in Supplemental Figure S4.

Supplemental Figure S6. Expression pattern of *Ol12G001732* and *OlGRXs*. **Supplemental Figure S7**. Expression of *Ol12G001732* in ramet shoots in response to exogenously supplied CEP1 peptides.

**Supplemental Table S1.** Genes up-regulated in the +N split treatment compared to the +N treatment

**Supplemental Table S2.** Genes up-regulated in the +N treatment compared to the -N treatment

**Supplemental Table S3.** Genes up-regulated in the +N treatment compared to the -N split treatment

**Supplemental Table S4.** Genes up-regulated in the -N treatment compared to the +N treatment

**Supplemental Table S5.** Genes up-regulated in the -N split treatment compared to the +N treatment

**Supplemental Table S6.** Genes up-regulated in the +N treatment compared to the +N split treatment

**Supplemental Table S7.** Cytokinin concentrations in the roots of +N, +N split, -N split, and -N treated ramets

**Supplemental Table S8.** Gene correspondence between *O. sativa* and *O. longistaminata*

**Supplemental Table S9.** Primers used for RT-qPCR analysis

## Acknowledgments

We are grateful to Drs. Motoyuki Ashikari, Tomoyuki Furuta, Takatoshi Kiba, Fanny Bellegarde, Mimi Hashimoto, Shoji Segami (Nagoya University), and Drs. Shinjiro Yamaguchi and Kiyoshi Mashiguchi (Kyoto University) for their helpful support and discussion.

---

**Supplemental Figure S1.**
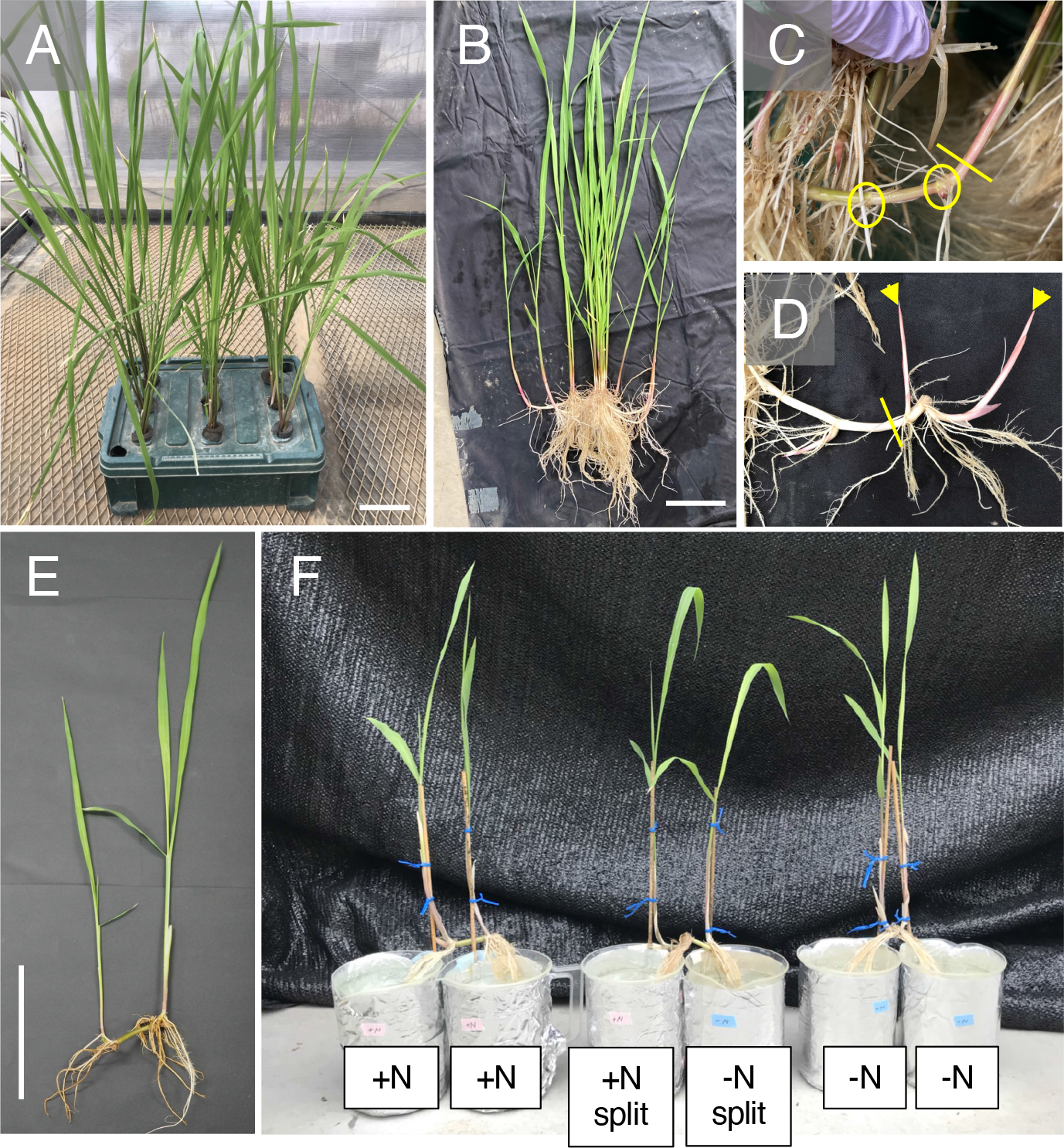
The split hydroponic experimental system for *O. longistaminata* ramet pairs. A, *O. longistaminata* hydroponic culture. The plants were maintained in a 10 L-container. Scale bar, 10 cm. B, A clonal colony of *O. longistaminata*. Scale bar, 10 cm. C, A rhizome grown in the hydroponic culture. Ovals identify adjacent nodes. The tip of a primary rhizome was cut at the yellow line to induce the outgrowth of secondary rhizomes. D, Growth of secondary rhizomes at adjacent nodes (arrowheads). The pair was excised from the parental colony at the yellow line for further growth. E, A ramet pair grown from adjacent rhizome nodes. Scale bar, 15 cm. F, Split treatments with different nitrogen conditions. Each pot contains 1 L of nutrient solution.

**Supplemental Figure S2.**
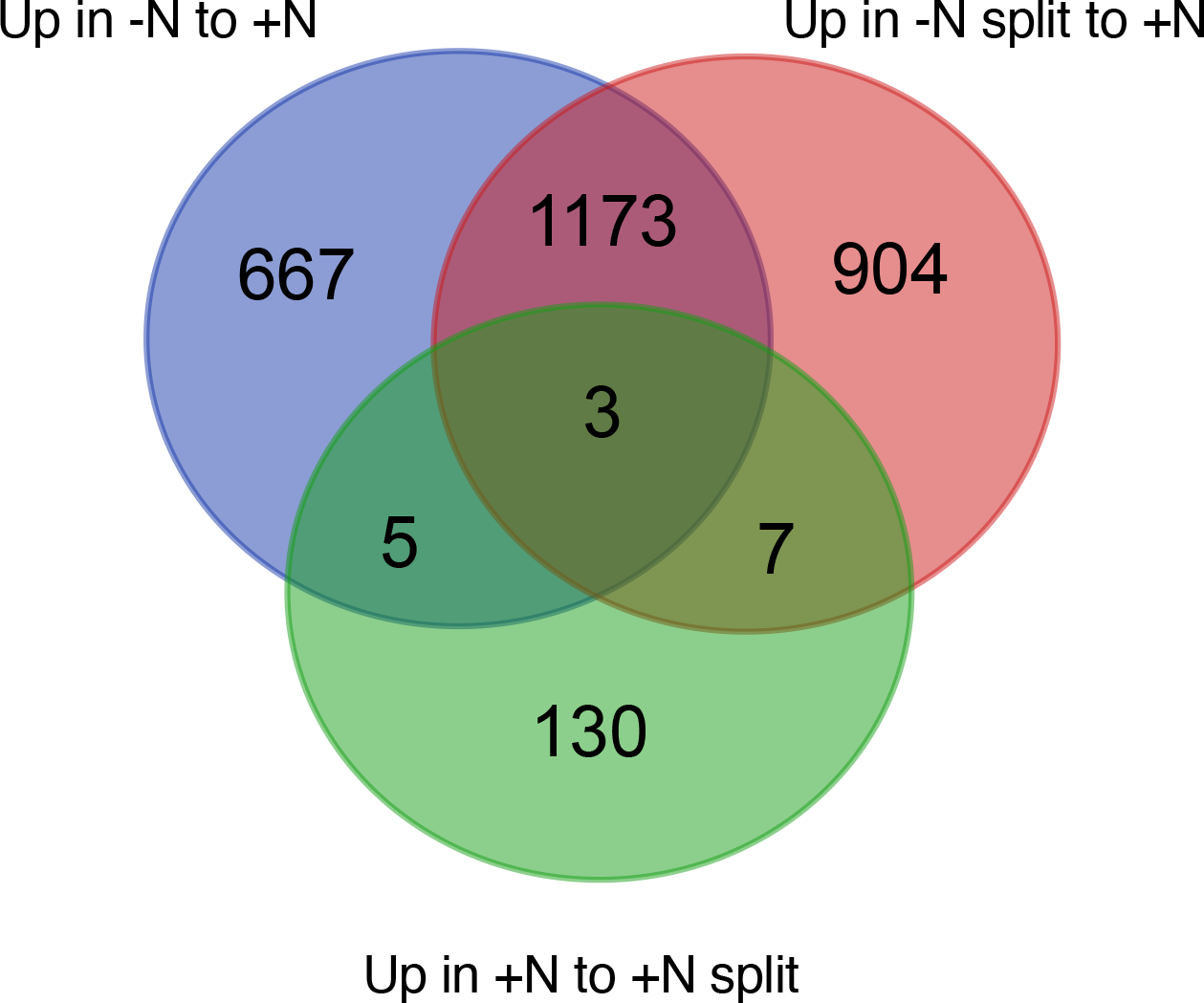
A Venn diagram showing the overlap among genes up-regulated in the -N treatment compared to the +N treatment (Up in -N to +N), those up-regulated in the -N split treatment compared to +N treatment (Up in -N split to +N), and those up-regulated in the +N treatment compared to +N split treatment (Up in +N to +N split).

**Supplemental Figure S3.**
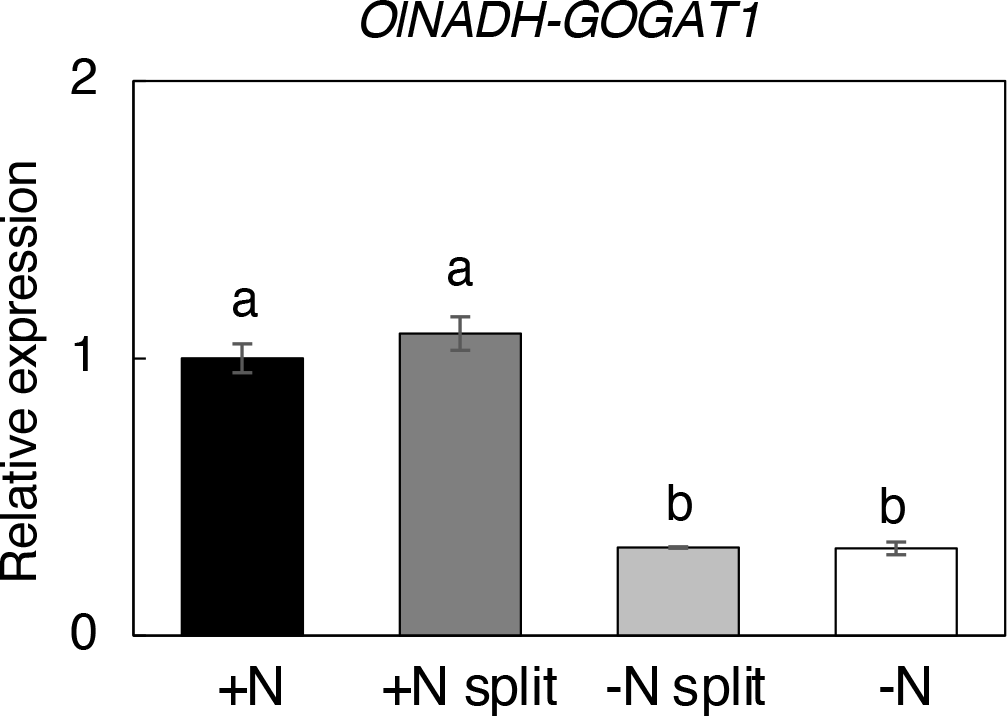
Transcript abundance of *OlNADH-GOGAT1* in the roots of ramet pairs after a 24-h split treatment as measured by RT-qPCR. The expression level of each gene, normalized by *TBC*, is expressed relative to that of the +N treatment defined as 1. Error bars represent SE of values for biological replicates (n = 3 or 4). Different lowercase letters at the top of each column denote statistically significant differences by Tukey’s honestly significant difference test (HSD) (*p* < 0.05).

**Supplemental Figure S4.**
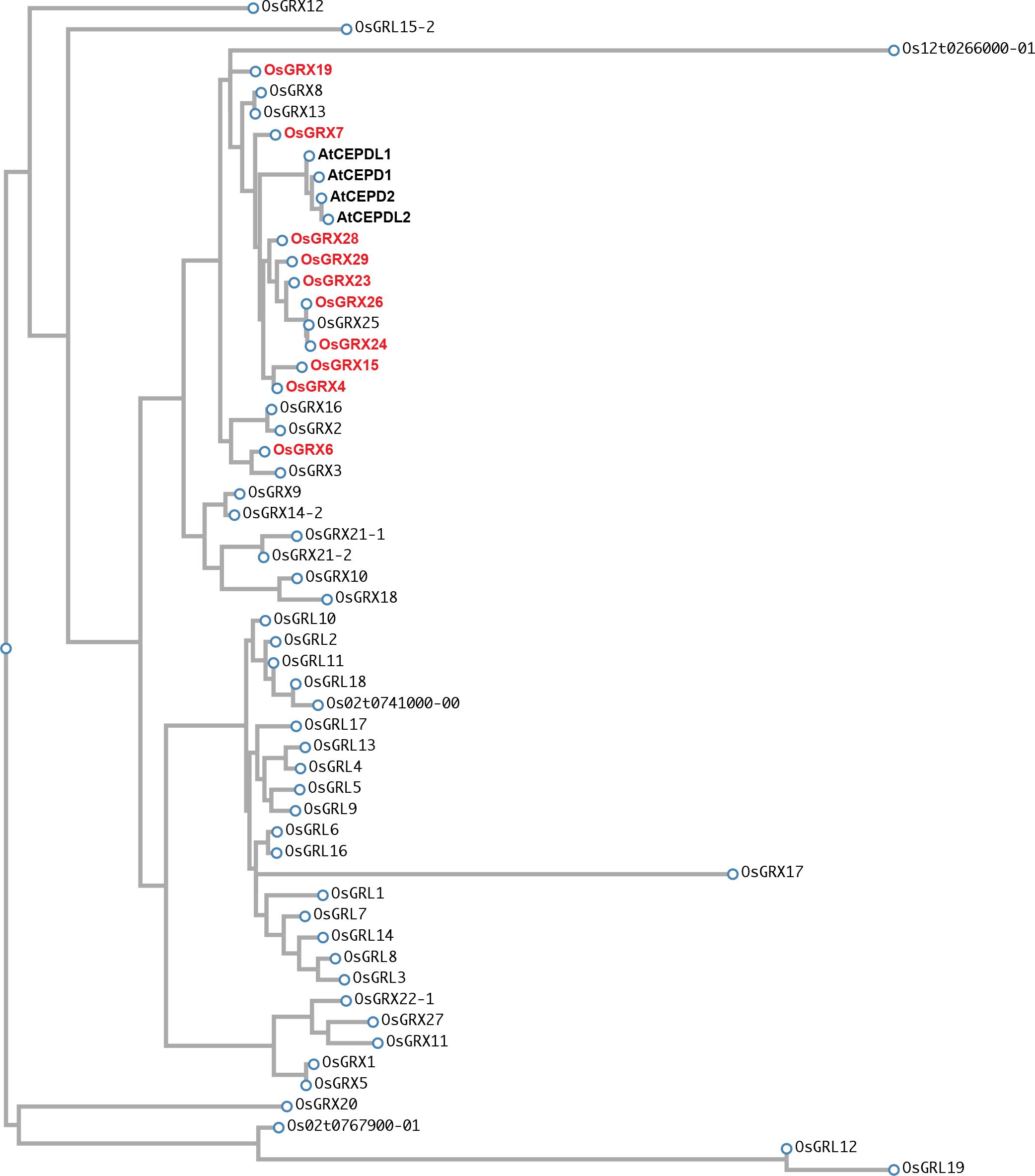
Phylogenetic analysis of Arabidopsis CEPDs (AtCEPD1, AtCEPD2, AtCEDL1, AtCEPDL2) and *O. sativa* glutaredoxin family proteins, GRX and GRL. Alignment and phylogenetic reconstructions were performed using the function “build” of ETE3 v3.1.1 (Huerta- Cepas et al., 2016) as implemented on the GenomeNet (https://www.genome.jp/tools/ete/). An alignment of the amino acid sequences used for this analysis is shown in Supplemental Figure S5. **Huerta-Cepas J, Serra F, Bork P** (2016) ETE 3: Reconstruction, Analysis, and Visualization of Phylogenomic Data. Mol Biol Evol **33**: 1635–1638

**Supplemental Figure S5.**
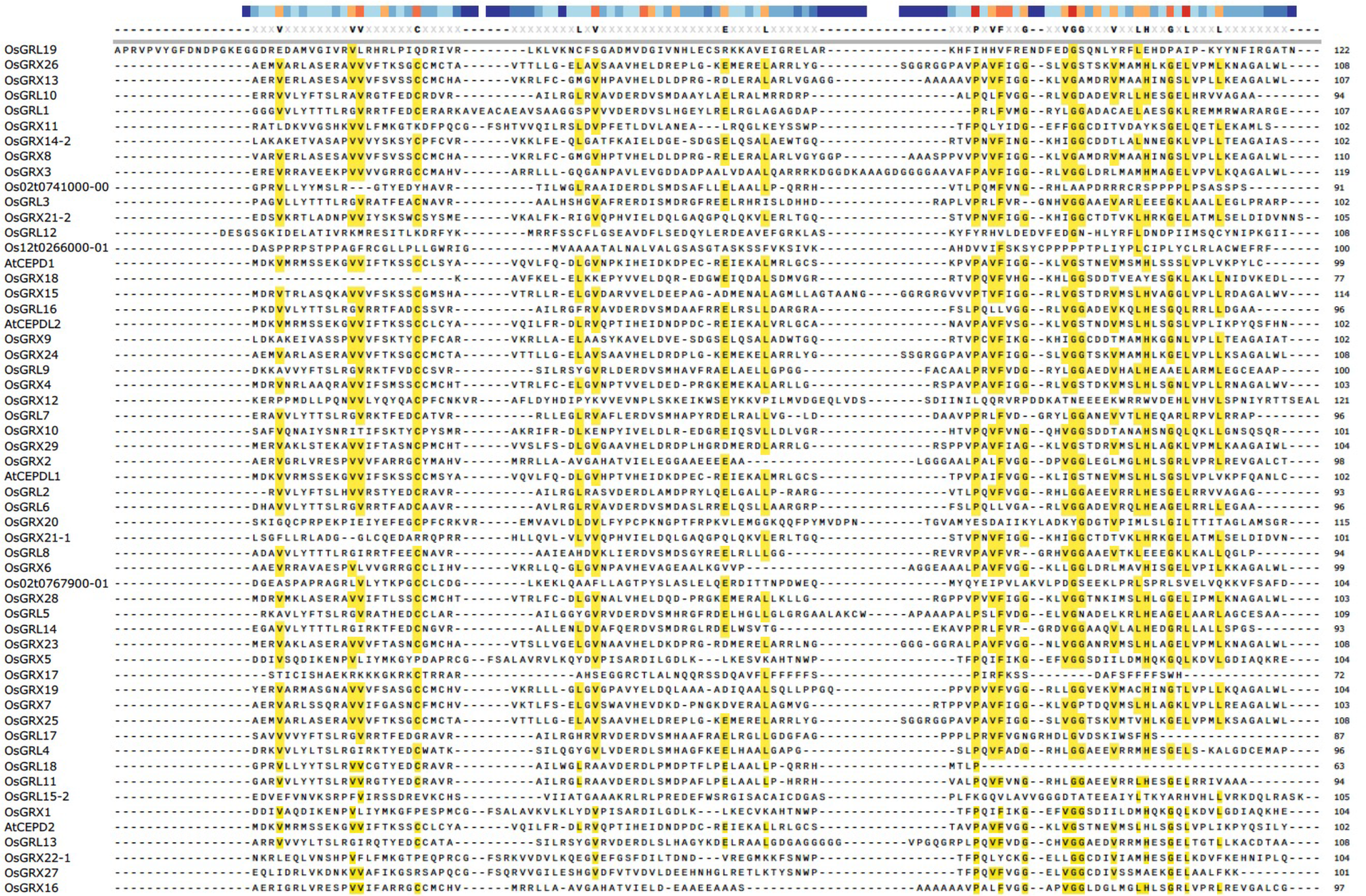
An alignment of the amino acid sequences used for the phylogenetic analysis in Supplemental Figure S4. AtCEPD1 (At1g06830), AtCEPD2 (At2g47880), AtCEPDL1 (At2g30540), AtCEPDL2 (At3g62960), OsGRX1 (Os01t0174900), OsGRX2 (Os01t0194600), OsGRX3 (Os01t0241400), OsGRX4 (Os01t0368900), OsGRX5 (Os01t0530400), OsGRX6 (Os01t0667900), OsGRX7 (Os01t0936000), OsGRX8 (Os02t0512400), OsGRX9 (Os02t0618100), OsGRX10 (Os02t0646400), OsGRX11 (Os03t0851200), OsGRX12 (Os04t0244400), OsGRX13 (Os04t0393500), OsGRX14 (Os04t0508300), OsGRX15 (Os05t0149950), OsGRX16 (Os05t0198200), OsGRX17 (Os05t0563900), OsGRX18 (Os06t0659500), OsGRX19 (Os07t0151100), OsGRX20 (Os08t0558200), OsGRX21-1 (Os08t0565800-01), OsGRX21-2 (Os08t0565800-02), OsGRX22 (Os10t0500700), OsGRX23 (Os11t0655900), OsGRX24 (Os11t0656000), OsGRX25 (Os11t0656400), OsGRX26 (Os11t0656700), OsGRX27 (Os12t0175500), OsGRX28 (Os12t0538600), OsGRX29 (Os12t0538700), OsGRL1 (Os01t0235900), OsGRL2 (Os01t0829400), OsGRL3 (Os02t0102000), OsGRL4 (Os02t0748800), OsGRL5 (Os03t0170800), OsGRL6 (Os03t0356400), OsGRL7 (Os03t0648800), OsGRL8 (Os04t0412800), OsGRL9 (Os04t0641300), OsGRL10 (Os05t0353600), OsGRL11 (Os05t0471350), OsGRL12 (Os06t0224200), OsGRL13 (Os06t0226100), OsGRL14 (Os07t0159900), OsGRL15 (Os07t0657900), OsGRL16 (Os07t0659900), OsGRL17 (Os08t0171333), OsGRL18 (Os08t0554699), OsGRL19 (Os10t0482900), Os02t0741000-00, Os02t0767900-01, Os12t0266000-01. Highly conserved amino acid residues are highlighted in yellow.

**Supplemental Figure S6.**
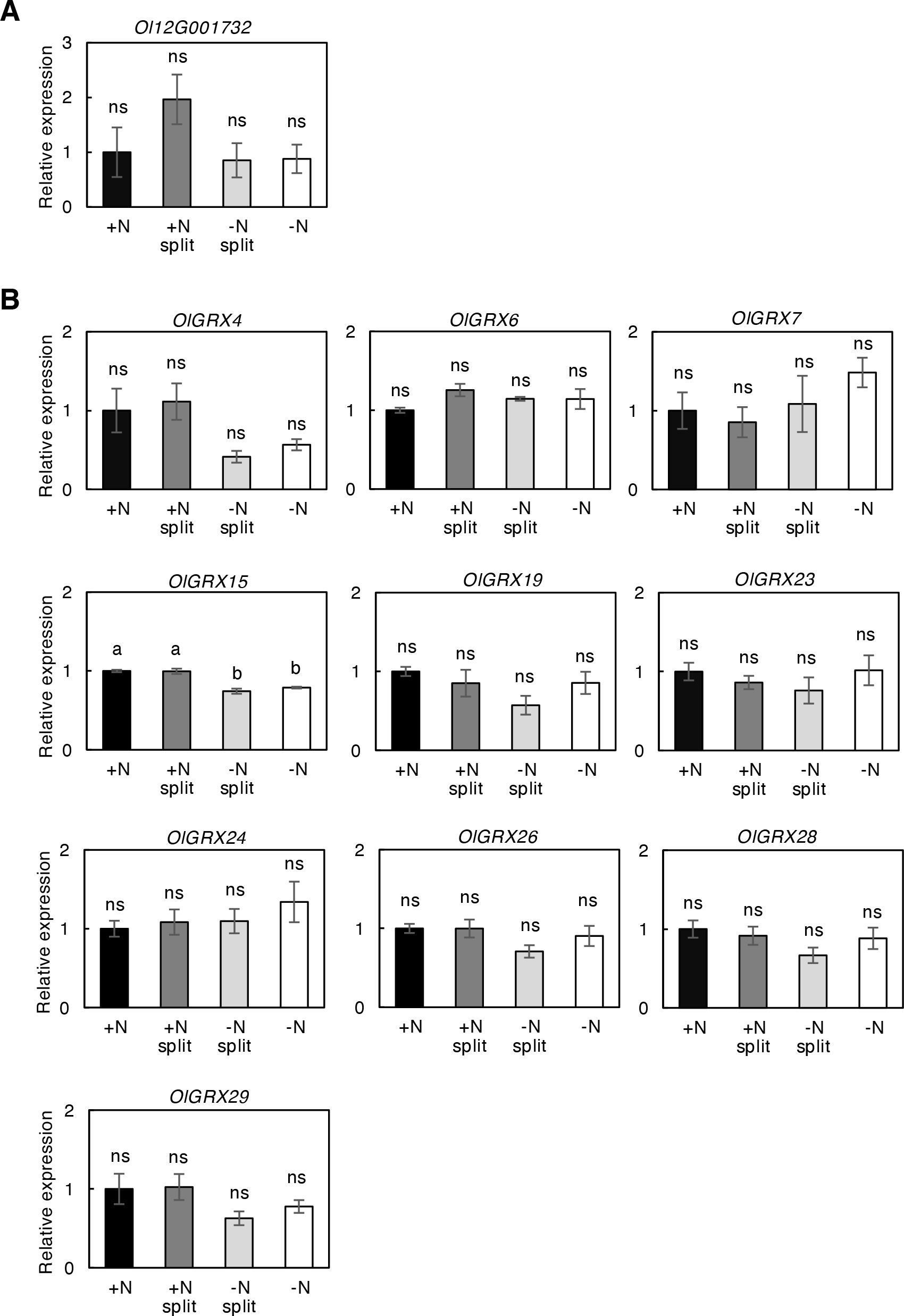
Expression patterns of *Ol12G001732* and *OlGRXs*. A, Expression pattern of *Ol12G001732*, the closest homolog of Arabidopsis *CEPR1.* B, Expression pattern of *OlGRXs*, homologs of Arabidopsis *CEPDs*, in the shoots of ramet pairs after a 24-h split treatment. The expression level of each gene normalized by *TBC* is expressed relative to that of the +N treatment defined as 1. Error bars represent the SE of values for biological replicates (n = 3 or 4). Different lowercase letters at the top of each column denote statistically significant differences by Tukey’s honestly significant difference test (HSD) (*p* < 0.05). ns, not significant.

**Supplemental Figure S7.**
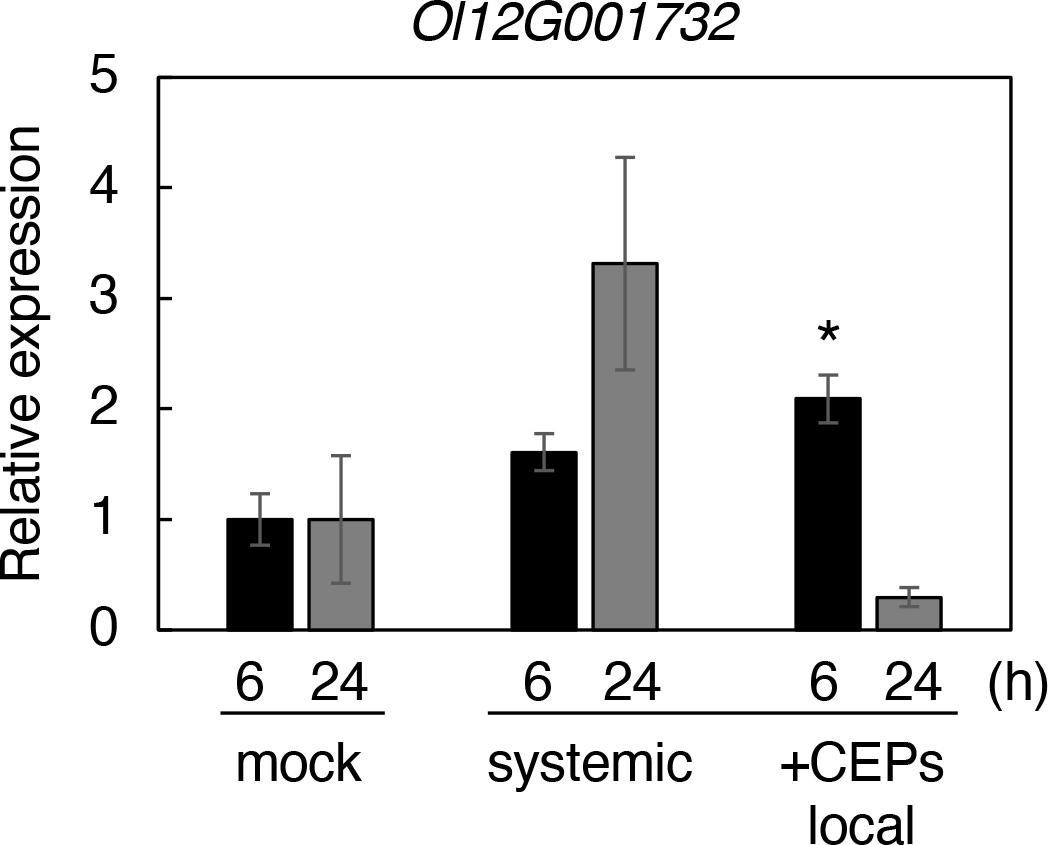
Expression of *Ol12G001732* in ramet shoots in response to exogenously supplied CEP1 peptides. The CEP1 peptide mix was applied to the roots of one ramet pair. Gene expression was measured in shoots from the local (+CEP local) and systemic side (systemic) after 6 and 24 h treatment. The expression level of *Ol12G001732*, normalized by *TBC*, is expressed relative to that of the +N mock treatment defined as 1. Error bars represent the SE of values for biological replicates (n = 3 or 4). \**p* < 0.05 (Student’s t-test) compared to the corresponding mock treatment.

